# Democratizing access to microfluidics: Rapid prototyping of open microchannels with low-cost LCD 3D printers

**DOI:** 10.1101/2023.09.26.558021

**Authors:** Kelsey M. Leong, Aileen Y. Sun, Mindy L. Quach, Carrie H. Lin, Cosette A. Craig, Felix Guo, Timothy R. Robinson, Megan Chang, Ayokunle O. Olanrewaju

## Abstract

Microfluidics offer user-friendly liquid handling for a range of biochemical applications. 3D printing microfluidics is rapid and cost-effective compared to conventional cleanroom fabrication. Typically, microfluidics are 3D printed using digital light projection (DLP) stereolithography (SLA), but many models in use are expensive (≥$10,000 USD), limiting widespread use. Recent liquid crystal display (LCD) technology advancements have provided inexpensive (<$500) SLA 3D printers with sufficient pixel resolution for microfluidic applications. However, there are only a few demonstrations of microfluidic fabrication, limited validation of print fidelity, and no direct comparisons between LCD and DLP printers. We compared a 40 µm pixel resolution DLP printer (∼$18,000 USD) with a 34.4 µm (<$380) LCD-SLA printer. Consistent with prior work, we observed linear trends between designed and measured channel widths ≥ 4 pixels on both printers, so we calculated accuracy above this size threshold. Using a standard IPA wash resin and optimized parameters for each printer, the average error between designed and measured widths was 2.11 ± 1.26% with the DLP printer and 15.4 ± 2.57% with the 34.4 µm LCD printer. The average coefficient of variation [CV] was ∼2% for both printers. Printing with optimized conditions for a low-cost water wash resin designed for LCD-SLA printers resulted in an average error of 2.53 ± 0.94% with the 34.4 µm LCD printer and 5.35 ± 4.49% with a 22 µm LCD printer. We characterized additional parameters including surface roughness, channel perpendicularity, and light intensity uniformity, and as an application of LCD-printed devices, we demonstrated consistent flow rates in capillaric circuits for self-regulated and self-powered delivery of multiple liquids. In conclusion, LCD printers are an inexpensive alternative for fabricating microfluidics, with minimal differences in fidelity and accuracy compared with a 20X more expensive DLP printer.

## Introduction

Microfluidic devices enable miniaturized and automated liquid handling across a wide range of applications ranging from diagnostics^1–3^ to electronics.^4^ Despite their potential to democratize access to automated fluid handling in a variety of settings, the high cost and centralization of microfabrication technology have limited the widespread deployment of microfluidics. The history of microfluidic fabrication has been thoroughly reviewed elsewhere,^5–7^ highlighting the progression from expensive and time-consuming methods, such as cleanroom fabrication, towards more economical and rapid prototyping methods like laser cutting and 3D printing. The swift and frugal prototyping offered by 3D printing, as well as the ease of realizing features with complex 3-dimensional structures, accelerates discovery and enables new design paradigms in microfluidics.

Prior work^8^ compared different 3D printing technologies – including fused deposition modeling (FDM), stereolithography (SLA), and polyjet – with the goal of achieving rapid, cost-effective, and high-fidelity microfluidic devices. SLA-based 3D printing yielded high-fidelity and high-resolution features for microfluidic applications and consequently digital light processing (DLP) using high-resolution projectors has become the gold standard^9–11^ for 3D printing microfluidics. Custom 3D-printers for microfluidic applications have been developed and microchannels with cross sections as small as 18 µm X 20 µm have been reported.^12^ However, the initial acquisition costs of high-resolution DLP printers are prohibitive (typically ≥ $10,000), which precludes access to many researchers who would benefit from rapid and inexpensive prototyping.

Recent advances in 3D printing technology and liquid crystal displays (LCDs) have provided access to SLA printers with high resolution and low cost. LCD SLA printers uses LCD screens instead of projectors to modulate illumination of photosensitive resin **(Supplementary Figure 1a)**.^13,14^ Over the past decade, advancements in LCD technology spearheaded by the inclination to improve the resolution of smartphone, tablet, and computer displays while maintaining consumer affordability, have been adapted for LCD printer screens.^15^ Whereas LCD printing has traditionally been relegated to hobbyists requiring lower cost, smaller footprint, but not necessarily high-resolution printers, these recent improvements have leveled the playing field in print quality between LCD and DLP printers. Although there are reports of fabrication of positive-feature molds for PDMS microfluidic devices^16–18^ and direct 3D printing of microstructures;^19–23^ there is limited characterization of the effects of printing parameters like UV light intensity, exposure time, layer height, and light intensity uniformity across the build plate on microchannel fidelity and dimensional accuracy with LCD-SLA printers. For instance, another recent publication reported high-resolution 3D-printing of microfluidics using LCD printers^24^ focusing on resin development and versatility in applications. However, there was no direct comparison with “gold standard” DLP to highlight the design considerations and trade-offs that may arise when using less expensive LCD printers.

In this manuscript, we compare the microfluidic fabrication capabilities of two inexpensive (<$500 USD) LCD printers and a DLP printer (∼$18,000 USD). We present an in-depth characterization of low-cost LCD 3D printers for open-channel microfluidic device fabrication. We compare the measured and designed XY and Z dimensions of open microchannels in a standardized calibration microchip for the LCD and DLP printers. We also 3D print microchannels using a low-cost resin that can be washed with water rather than isopropanol, decreasing chemical/hazardous waste disposal concerns. Furthermore, we characterize channel perpendicularity, surface roughness, and feature reproducibility across the build plate. We also demonstrate an application of LCD printing by microfabricating a capillaric circuit for self-powered and self-regulated delivery of multiple liquids without external instruments or operator intervention.

## Materials and Methods

### Equipment & Software

We compared the performance of a commercially available DLP printer (CADWorks Profluidics 285D) and two LCD printers (Anycubic Mono X 6K, Phrozen Sonic Mini 8K Resin 3D printer). Microchannels and microfluidic circuits were designed using Solidworks (Waltham, Massachusetts, USA) and exported as Standard Tessellation Language (STL) files prior to 3D printing. The following printer and STL slicer combinations were used: Profluidics 285D printer and the CADworks Utility 6.44 slicer (CADworks3D, Toronto, Ontario, Canada), the Anycubic Photon Mono X 6K and the Anycubic V2.1.29 slicer (Anycubic, Shenzhen, Guangdong, China), and the Phrozen Sonic Mini 8K Resin 3D printer (Phrozen, Hsinchu, T’ai-wan, Taiwan) and Chitubox Basic 1.9.5 slicer (Chitubox, Shenzhen, Guangdong, China). Capillaric circuits were plasma treated with the Plasma Etch PE-25 (Carson City, Nevada, USA) at 56% power for 3 minutes to achieve a hydrophilic surface for capillary-driven flow. Image capture, sidewall angle, XY measurements **(Supplementary Figure 2)**, and height measurements were performed with the Keyence VH-S30B digital microscope (Itasca, Illinois, USA). Image capture and height measurements were performed using the FEI XL30 Sirion scanning electron microscope (SEM) (Hillsboro, Oregon, USA). Surface roughness measurements were taken with the Keyence VR-5000 Wide-area 3D Measurement System (Itasca, Illinois, USA). Videos of liquid delivery in the microfluidic chips were taken using an iPhone XS (Apple, Cupertino, CA, USA).

## Materials

The following commercially available resins were used in the printer optimization experiments: Clear Microfluidics Resin V7.0a (CADworks3D, Toronto, ON, Canada) and Water-Wash Resin+ (AnyCubic, Shenzhen, Guangdong, China). Isopropanol was sourced from Sigma-Aldrich (Burlington, Massachusetts, USA), and colored food dyes (Watkins, Winona, Minnesota, USA) were employed for flow characterization in 3D printed capillaric circuits.

### Evaluation of dimensional accuracy using calibration chips

Two different sets of calibration chips with microchannels of varying widths and heights were fabricated to test the 3D printers’ XY and Z resolution limitations **(Supplementary Figure 1B)**. We developed calibration chips to determine the optimal layer thickness for the prints. We designed 3 height calibration chips (10 µm, 20 µm, and 30 µm layer thickness) for each printer with channels at a uniform XY size of 10 pixels. The channel heights began at 1 layer and peaked at the number of layers that did not result in a total height > 160 µm. XY calibration chips made for the Anycubic Mono X 6K printer and Phrozen Sonic Mini 8K were designed to test the manufacturers’ specified pixel resolution of 34.4 µm^25^ and 22 µm^26^, respectively. The channels on the calibration chips were aligned with the LCD printer’s pixel grid **(Supplementary Figure 1A)** to avoid errors associated with features that fell between gridlines and could not be sliced or printed properly with the LCD printer. To test XY resolution, microchannel widths ranged from 1-10 pixels increasing in whole pixel steps at a uniform Z-height of 160 µm.

The calibration chip designs were exported as STL files, converted into G-code with the slicer, and 3D printed. The chips printed with the CADworks resin were washed in an isopropanol (IPA) bath and dried using a compressed nitrogen gun (Cleanroom World, Centennial, Colorado, USA). In contrast, chips printed with the Anycubic Water-Wash Resin+ were washed in a deionized water bath and dried using a compressed nitrogen gun. The washing and drying steps were repeated, as needed, for both resins to ensure that any remaining uncured resin was removed from the part. After drying, the microchips were placed in a Formcure (Formlabs, Somerville, Massachusetts, USA) to post-cure for 30s on each side of the chip, for a total of 1 minute.

### Capillaric circuits for pre-programmed instrument-free liquid delivery

Capillaric circuits were fabricated to test the performance of the 3D printed parts and showcase a variety of self-powered fluidic elements that enable hands-free and minimally-instrumented liquid delivery. Once printed, the chips were washed, dried, and post-cured as described above. After post-curing, the chips were plasma-treated and sealed with hydrophobic tape (ARseal™ 90697, Adhesives Research, Glen Rock, Pennsylvania, USA). Food coloring (13% v/v in water) was used to test flow in the capillaric circuits. Non-woven cleanroom paper (AAWipes, Ann Arbor, Michigan, USA) was used as the capillary pump and fastened to the capillaric circuit using a rubber band.

## Results and Discussion

The specifications for the DLP and LCD printers used in this study are summarized in **Table 1**. The CADworks DLD printer has a native (projector) pixel size of 40 µm and a dynamic (software-enabled) pixel size of 28.5 µm. Meanwhile, the Phrozen and Anycubic LCD-SLA printers had pixel sizes of 22 µm and 34.4 µm, respectively. The LCD printers were smaller, lighter, and much (≥40X) less expensive than the DLP printer. The LCD printers operate at 405 nm, while the DLP printer operates at 385 nm.

**Table 1:** Comparison of specifications for the DLP and LCD printer.

|  | 3D Printer |  |  |
| --- | --- | --- | --- |
|  | CADworks Profluidics 285D | Anycubic Mono X 6K | Phrozen Sonic Mini 8K |
| Printing method | Digital Light Processing (DLP) stereolithography | LCD or Masked Stereolithography | LCD or Masked Stereolithography |
| Pixel Resolution | 2,750 x 1,550 pixels | 5,760 x 3,600 pixels | 7,536 x 3,240 pixels |
| Pixel size | 40 $\mu\text{m}$ (native)<br>28.5 $\mu\text{m}$ (dynamic) | 34.4 $\mu\text{m}$ | 22 $\mu\text{m}$ |
| Build area | 110 x 62 x 120 | 245 x 197 x 122 | 165 x 72 x 180 |
| **XYZ (mm)** |  |  |  |
| **Operating wavelength** | 385 nm | 405 nm | 405 nm |
| **Machine footprint XYZ (mm)** | 500 x 570 x 590 | 475 x 290 x 270 | 290 x 290 x 430 |
| **Light intensity** | 5 – 6 mW/cm<sup>2</sup> | 5 – 6.5 mW/cm<sup>2</sup> | 2.8 mW/cm<sup>2</sup> |
| **Instrument Weight** | 56 kg | 11 kg | 13 kg |
| **Cost (USD)** | \$17,995* | \$382* | \$449.99* |

\*Documented price as of 9/8/23

### Comparison of channel widths and depths between DLP and LCD printers

LCD printing parameters such as exposure time, off-time, and UV power were optimized to produce high-resolution and high-fidelity features. **Supplementary Table 1 and Supplementary Figure 3** summarize these parameters and describe the optimization process. We printed the same XY calibration chips with CADworks resin (IPA wash resin) on both the CADworks Profluidics (DLP) **(Figure 1A)** and Anycubic Mono X 6K (34.4 µm LCD printer) **(Figure 1B)**. We printed open microchannels at different layer thicknesses (10 µm, 20 µm, or 30 µm) to investigate the tradeoff between channel fidelity and print time. 20 µm layer heights minimized sidewall channel roughness while also maintaining an overall print time below 40 minutes **(Supplementary Figure 4)**. SEM photos provide qualitative comparisons between microchannels produced by the printers. For XY feature comparisons, we fixed microchannel depth at 160 µm (13 layers) and systematically varied channel widths from 1 to 10 pixels. The 1-pixel channels were poorly defined on both printers. The 2- and 3-pixel channels were better defined, although they lost dimensional integrity near channel edges and resembled comet or brush stroke shapes, more noticeable on the channels printed by the LCD printer (**Figure 1B-ii**). Besides slight rounding at channel corners, features above 3 pixels did not have noticeable defects. This is similar to prior work with DLP printers where minimum microchannel widths were 2.5 – 4X the printer’s minimum pixel size;^10^ in keeping with this trend we analyzed the accuracy of channel widths ≥ 4 pixels.

**Figure 1:**
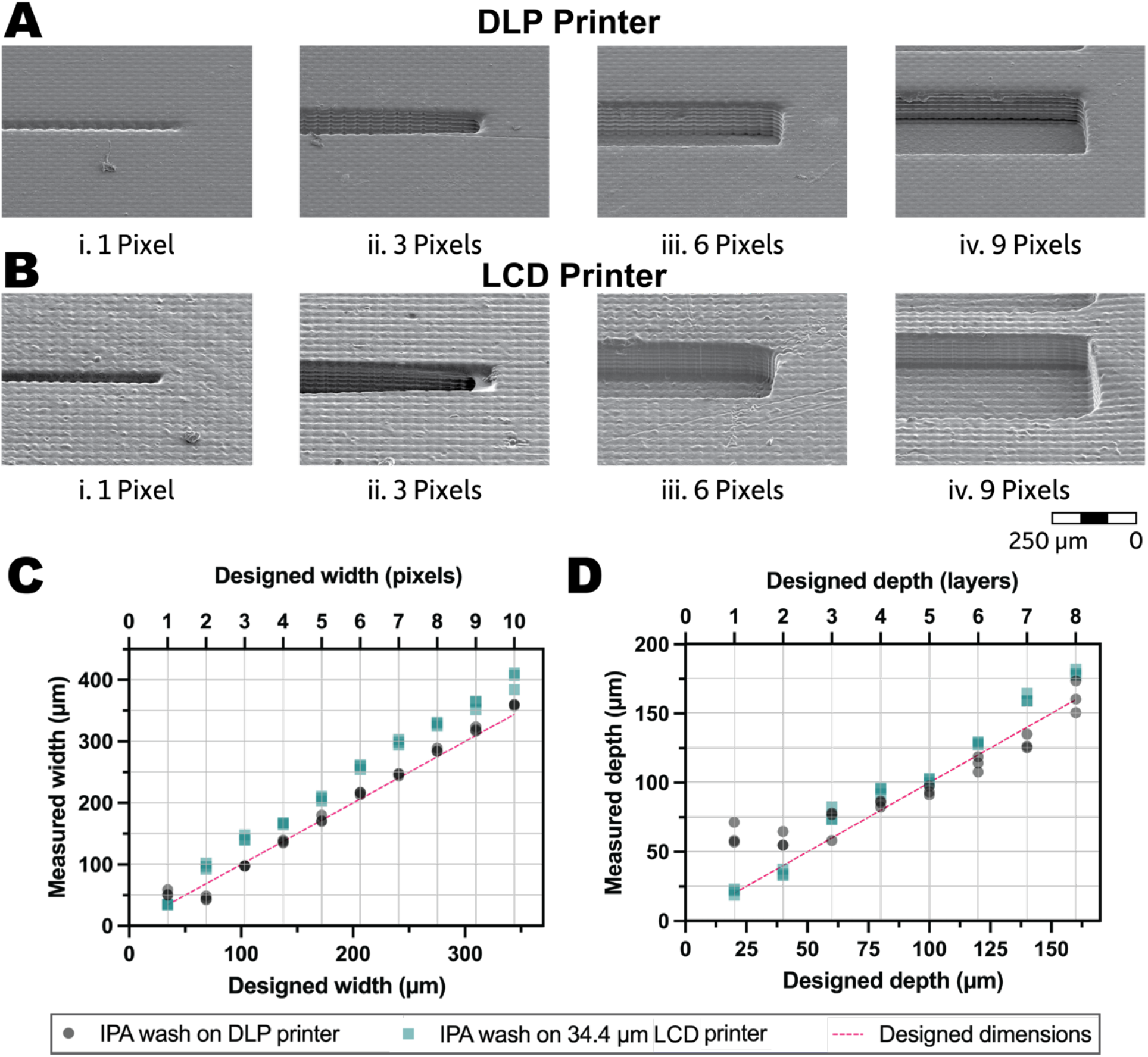
Comparison between microchannels printed with the same resin on LCD and DLP printers. **(A)** SEM images of microchannels printed at a uniform height with XY widths varying from 1 pixel (34.4 µm) to 9 pixels (309.6 µm). Microchannels printed on the DLP printer with IPA wash resin. Channels 3 pixels (103.2 µm) and below had difficulty maintaining a linear shape, but this was not prevalent above 3 pixels. **(B)** Microchannels printed on the 34.4 µm LCD printer with IPA wash resin show slight rounding of edges and “cometing” effect of channels below 3 pixels (103.2 µm), but otherwise no obvious defects. **(C)** Microchannels printed with the DLP had excellent agreement between designed and measured widths, while channels from the LCD printer were larger but maintained a good linear trend as pixel size increased. **(D)** The LCD printer produced depths that had a linear trend with the designed dimensions at all layer sizes, while the DLP printer maintained a linear trend for channels 3 layers (60 µm deep) and above.

Microchannel widths from the DLP printer **(Figure 1C)** strongly correlated with designed dimensions above 4 pixels (137.6 µm) with an average error of 2.11 ± 1.26% for channels between 4 and 10 pixels (344 µm). Channels printed with the DLP printer were very precise with average coefficient of variation [CV] of 2.17 ± 2.53% across three replicate channels on the same calibration chip. While the calibration chip was designed for the Anycubic LCD’s 34.4 µm pixel size, these results demonstrate that the DLP printer was still able to produce accurate feature sizes due to its dynamic printing capabilities.

Microchannels printed with the LCD printer were larger than those obtained with the DLP printer (and the designed dimensions) **(Figure 1C)**. The average difference between measured and designed dimensions for the LCD printer was 15.4 ± 2.57% for features ≥ 4 pixels (137.6 µm). Nevertheless, replicated microchannels (N=3) on the LCD printer were very precise for each pixel size with an average [CV] = 1.8 ± 1.11%. XY measurements from the LCD printer also showed consistent linearity for features ≥ 4 pixels.

The LCD printer had a similar depth profile to the DLP printer **(Figure 1D)**. There was good linearity and lower variation between designed and measured dimensions across all layers. The deviation between measured and designed depths with the LCD printer was on average 12.66 ± 8.01%, compared to 12.91 ± 15.37% with the DLP printer (omitting the 20 µm deep channel for the DLP). For the LCD printer, there was minimal variation between depths of triplicate microchannels printed on the same pixel calibration chip with an average [CV] = 4.01 ± 3.56% compared to [CV] = 2.64 ± 3.22 of channels produced by the DLP printer. Taken together, our width and depth measurements indicate that the LCD printer can fabricate microchannels that are suitable for many microfluidic applications.

### Comparison of channel perpendicularity and surface roughness between DLP and LCD printers

To investigate microchannel perpendicularity especially for small features we printed channels with four different aspect ratios on the DLP and LCD printers **(Figure 2A)**. We chose aspect ratios that we would naturally come across when designing microfluidic features; however, we specified low height (40 µm) and width (103.2 µm) values to push the limits of the LCD printers. The DLP printer produced precise channels on target with the designed sidewall angles of 90° across all aspect ratios **(Figure 2A-i)**, although the 2:1 aspect ratio channel was much shallower than expected. The LCD printer generated channels within ±10° of the designed angle at the two higher aspect ratios **(Figure 2A-ii)** while the two lower aspect ratio channels had much wider side wall angles than designed.

**Figure 2:**
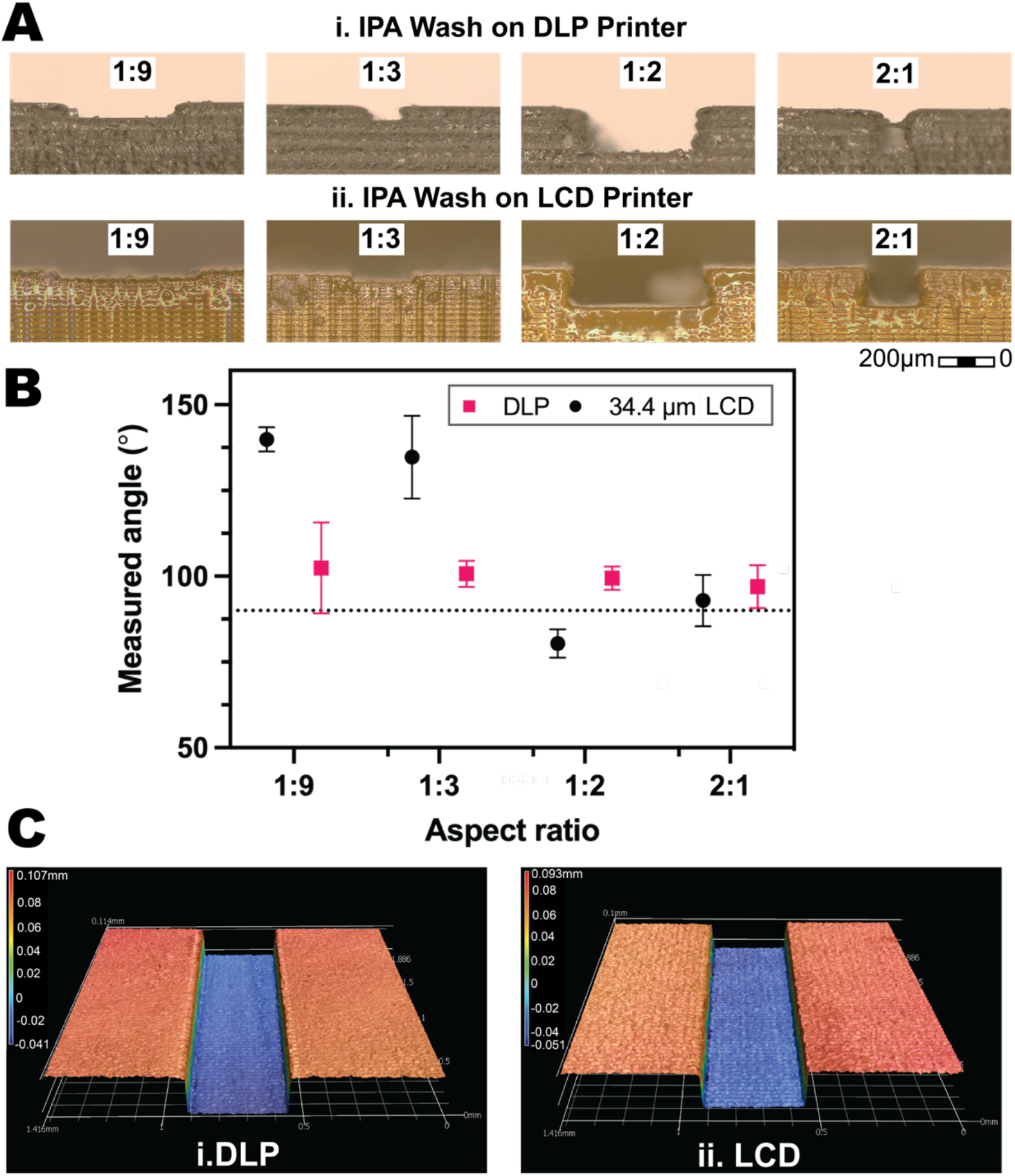
Evaluation of channel perpendicularity and surface roughness between features printed with the same resin on LCD and DLP printers. **(A)** Channels with various aspect ratios were printed on the DLP and LCD printers to investigate how channel design influences perpendicularity. **(b)** The DLP printer produced channels with measured angles in close agreement with the designed 90° (dashed line). The LCD printer obtained results closer to the designed architecture when the aspect ratio was higher. **(c)** 3D surface profile images of channels printed on the i) DLP and ii) LCD printers show similar surface roughness.

Surface roughness increases flow resistance, which can cause the experimental flow rates to diverge from the predicted flow rate.^27^ We quantified the surface roughness of 3D printed channels from both the DLP and LCD printer. We obtained 3D scans of 10-pixel wide microchannels from 3 different chips using a Keyence optical profilometer **(Figure 2C)**. We randomly selected 400-pixel x 100-pixel areas (N=9) inside the channels **(blue regions in Figure 2C)**. Printed features had average surface roughness of 1.52 ± 0.54 µm and 2.09 ± 0.44 µm on the LCD and DLP printers, respectively, with the surface texture from the LCD-printed parts being slightly more prominent inside and outside the channels.

### 3D printing with low-cost, water wash resin on two LCD printers

To demonstrate that we could optimize the print conditions of multiple LCD printers from different manufacturers, we designed and printed a pixel calibration chip on a second LCD printer – the Phrozen Sonic Mini 8k with 22 µm pixel resolution, which was released during our experimentation with the first LCD printer. We initially printed calibration chips with the IPA-wash Clear Microfluidics Resin V7.0a (CADworks3D, Toronto, Ontario) that cost $510/kg and showed no significant differences in print fidelity **(Supplementary Figure 5)**. Subsequently, we ran additional experiments to test the LCD printers with a resin specifically formatted for its 405 nm light engine (Water Wash+, AnyCubic, Shenzhen, Guangdong, China). As an added benefit, the water-washing capabilities reduce chemical/hazardous waste disposal concerns, and the lower price ($34/kg) decreases operating costs.

While the goal of this work was to characterize printer hardware rather than different resin formulations, the ability to create high-fidelity, small features is a function of the printer hardware, the resin formulation, and printer-resin compatibility. Using the water-wash resin, the 34.4 µm LCD printer demonstrated an average difference between designed and measured widths of 2.53 ± 0.94% for microchannels ≥ 4 pixels and a coefficient of variation of 2.48 ± 1.43% (N=3) **(Figure 3A)**. Microchannel depths **(Figure 3B)** were larger than designed dimensions with an average difference of 18.68 ± 2.23% (omitting the first layer) but maintained an overall linear trend. Meanwhile, the 22 µm LCD printer with water-wash resin and microchannels ≥ 4 pixels, had an average difference 5.35 ± 4.49% between designed and measured widths with a coefficient of variation of 3.72 ± 3.02% **(Figure 3A)**. There was an overall linear trend between designed and measured widths for channels ≥ 4 pixels and microchannel depths were very close to the designed dimensions with an average difference of 4.04 ± 3.43% between measured and designed dimensions **(Figure 3B)**. These results suggest that we can obtain repeatable results on two different LCD printers for microchannels with sizes and aspect ratios relevant to microfluidic applications. Our approach may be broadly generalizable to other LCD printers with similar pixel resolution and price points.

**Figure 3:**
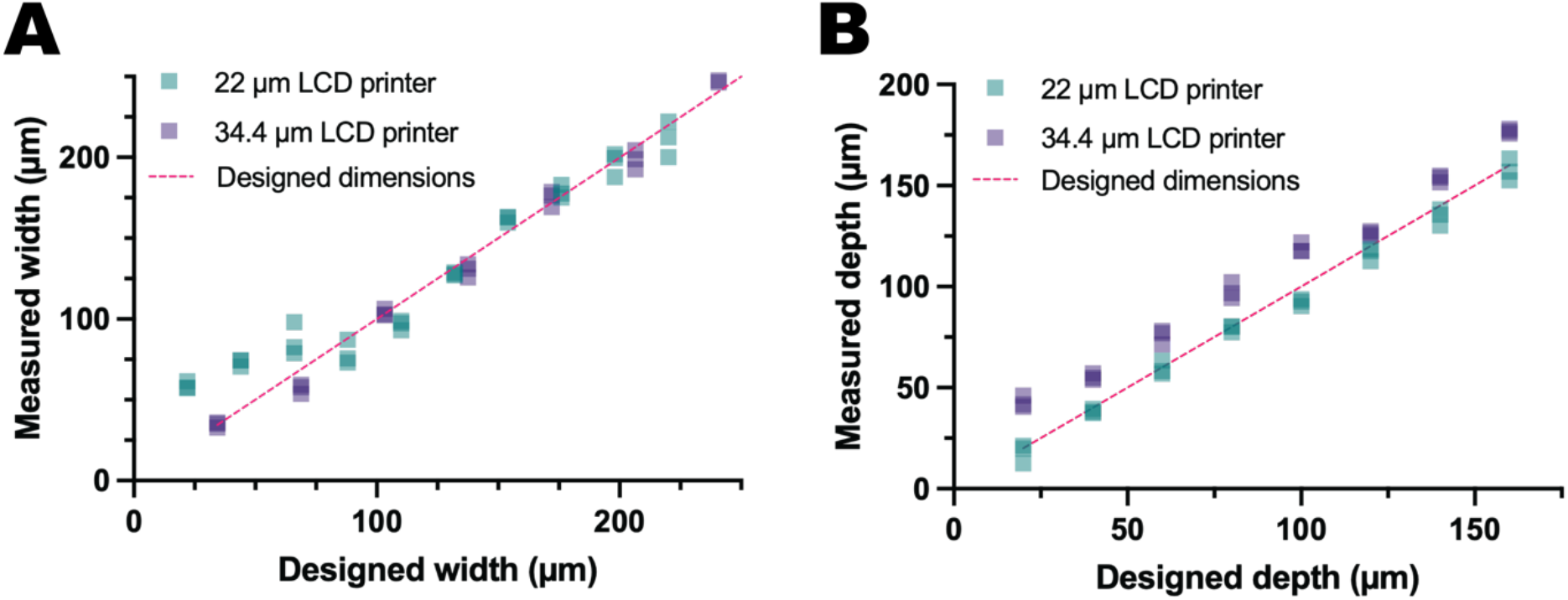
Microchannels printed with a low-cost, water wash resin on 34.4 µm-pixel and 22 µm-pixel LCD printers. **(A)** There was excellent agreement with designed and measured dimensions across all widths for the 34.4 µm LCD printer. Meanwhile, the 22 µm LCD printer produced microchannels that showed a linear trend with the designed dimensions ≥ 4 pixels (88 µm). **(B)** Microchannels printed on the 34.4 µm LCD printer were ∼18% deeper than expected across all layers while channels on the 22 µm LCD printer showed excellent agreement between designed and measured depths across the layer thicknesses tested (20 – 160 µm).

### Pre-programmed, instrument-free liquid delivery with capillaric circuits

We designed capillaric circuits composed of multiple fluidic elements **(Figure 4A)** that are powered by capillary forces to demonstrate the utility of microchannels fabricated with LCD printers. CAD designs were aligned to the pixel grid and printed using water wash resin on the 34.4 µm LCD printer. The smallest features in the capillaric circuit were the trigger valve, retention valve, and flow resistor, which had feature widths ranging from 5 - 6 pixels and depths of 200 µm. Other than slight narrowing of the flow resistor **(Figure 4A-iv)**, there were no obvious defects in capillaric circuits fabricated with the LCD printers. Three capillaric circuits were printed, plasma treated, and tested with food dye solutions to demonstrate automated, sequential liquid delivery performed on LCD-printed microchips.

**Figure 4:**
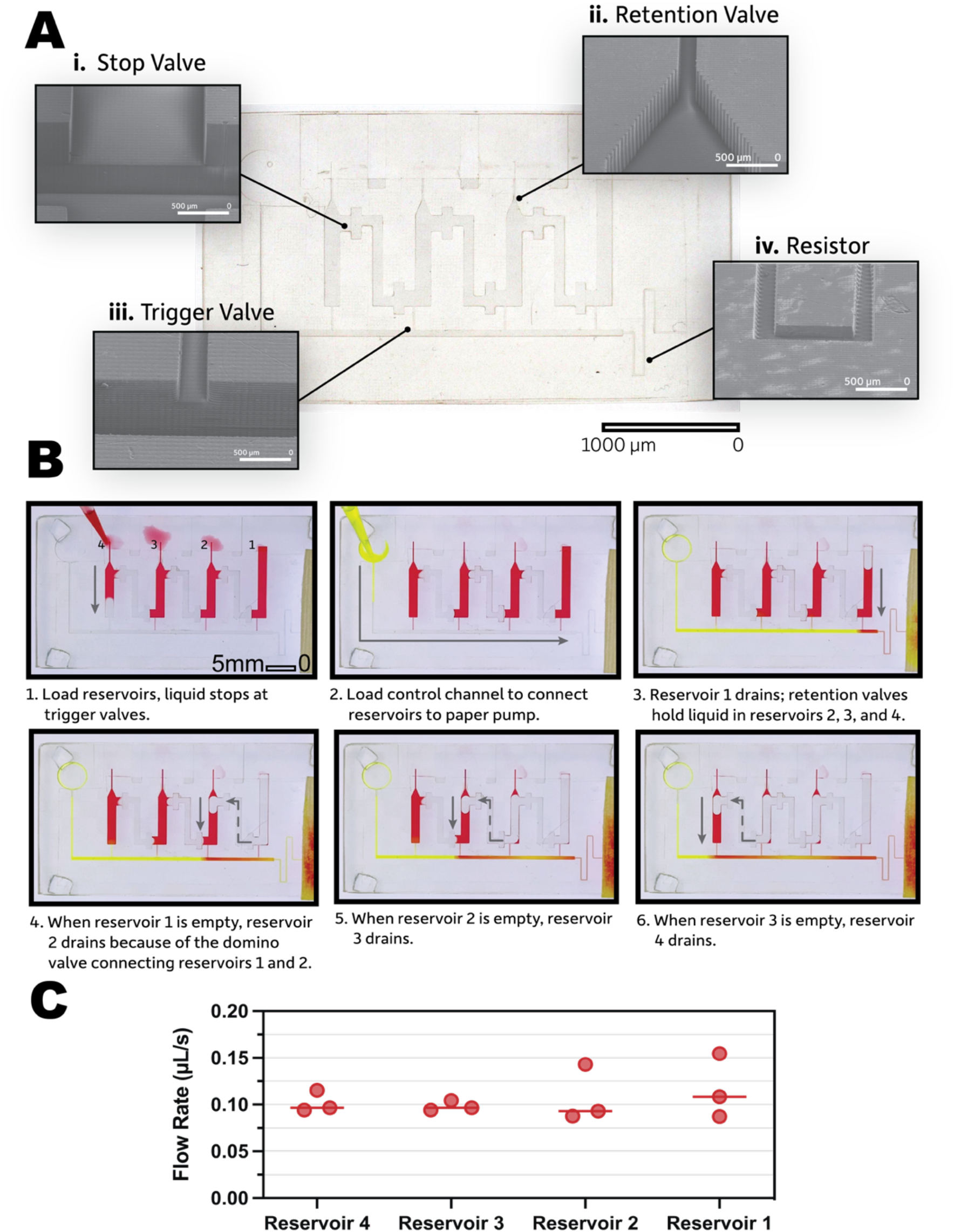
Pre-programmed sequential liquid delivery with capillaric circuits 3D-printed with low-cost, water-wash resin on LCD printer. **(A)** The 3D printed circuit is shown in the center and insets show SEM images of the fluidic components: (i) Stop valve, (ii) Retention valve, (iii) Trigger valve, and (iv) Resistor. Scale bars on SEM images are 500 µm. **(B)** Screenshots from supplementary video 1 showing pre-programmed sequential liquid delivery in a capillaric circuit. The screenshots progress from filling the reservoirs and control channel (1,2), through the sequential drainage of each reservoir (3 – 6). **(C)** Flow rates were consistent across all reservoirs and across three 3D-printed microchips.

Operation of capillaric circuits is documented in **Supplementary Movie 1** and **Figure 4B** and follows our previous work describing the operating principle of capillary domino valves.^29^ First, the user loads liquids into reservoirs where trigger valves hold liquid in place **(Figure 4B-1)** until pre-programmed sequential liquid delivery is initiated by loading the control channel that connects the reservoirs to the capillary pump **(Figure 4B-2)**. Unlike reservoirs 2, 3, and 4 that have retention valves preventing their immediate drainage, reservoir 1 immediately drains when connected to the capillary pump **(Figure 4B-3)**. When reservoir 1 is empty, the domino valve connecting reservoirs 1 and 2 enables drainage of reservoir 2 **(Figure 4B-4)**. This triggers a domino effect where each reservoir drains once the preceding reservoir is completely empty **(Figure 4B-5,6)**. We compared flow rates from each reservoir across 3 different microchips **(Figure 4C)** and found minimal differences when comparing the flow rates, demonstrating reproducible results from print to print. These results show that we can fabricate high-fidelity, functional capillaric circuits for hands-free and instrument-free liquid delivery using an inexpensive LCD printer.

If light is not properly collimated through the LCD display, the illumination across the build plate is non-uniform and results in imprecise features.^28^ We performed light intensity measurements to determine the uniformity of the light intensity across the build plate of the two LCD printers **(Supplementary Figure 6)** and measured features of capillaric circuits printed on different areas of the build plate **(Supplementary Figure 7)**. We found that the UV intensity was generally uniform for both the LCD printers and the measured widths of the capillaric features depicted high precision regardless of print location.

## Conclusion

This paper serves as both a successful validation and a printing guide for low-cost LCD printers. We demonstrate that a suitable trade-off can be found between 3D printer cost and microchannel fidelity and accuracy for open microchannels. We compared a 40-µm pixel DLP printer (∼$18,000) to a 34.4-µm pixel LCD printer (<$380) and obtained linear trends between designed and measured dimensions for both printers without significant differences in surface roughness and channel perpendicularity. We varied LCD printer parameters (e.g., exposure time, power intensity, layer height) to achieve high-fidelity microfluidic features and characterized microchannel properties (e.g., dimensional accuracy, surface roughness, perpendicularity) between DLP and LCD printers. We also fabricated microchannels using a low-cost ($34/liter) water wash resin on both the 34.4 µm-pixel LCD printer and a 22 µm-pixel LCD printer demonstrating the versatility of our optimization. This flexibility may allow researchers to test resins with different material properties, such as elasticity, hydrophilicity, optical clarity, or biocompatibility to broaden their functionality for biomedical applications.

Frugal 3D printing of microfluidics fosters more inclusive access to liquid handling solutions and allows a broader range of researchers to design and apply microfluidics to their research.

## Author Contributions

K.L. was involved in the conceptualization, methodology, investigation, validation, formal analysis, data curation, project administration, and completed the initial draft. A.S. and M.Q. participated in investigation, formal analysis, and data curation. C.C. and C.L. helped with visualization, paper editing, and software, and C.L. aided with investigation. F.G. assisted with conceptualization, investigation, and data curation. T.R. and M.C. contributed to paper edits and supervision. A.O. provided supervision and aided with methodology, funding acquisition, conceptualization, and paper editing.

## Acknowledgments

We are grateful to Ashleigh Theberge, Jodie Tokihiro, Yuting Zeng, Dan Ratner, Lucas Meza, for helpful comments, access to instruments, and support.

## Funding Information

We are grateful for funding from the Atlanta Center for Microsystems Engineered Point-of-Care Technologies and the National Science Foundation (Award #2223537). Part of this work was conducted at the Washington Nanofabrication Facility / Molecular Analysis Facility, a National Nanotechnology Coordinated Infrastructure (NNCI) site at the University of Washington with partial support from the National Science Foundation via awards NNCI-1542101 and NNCI-2025489. We acknowledge equipment support from the M.J. Murdock Diagnostics Foundry for Translational Research and the Molecular Analysis Facility at the University of Washington.

## Notes

https://doi.org/10.5281/zenodo.8347134

### Competing Interest Statement

The authors have declared no competing interest.

### Summary of Updates

We have completed the following changes to our manuscript, as suggested by reviewers, to prepare for this submission: (1) reduced the length of the abstract, (2) deemphasized the importance of capillary microfluidics and extended the purview of our paper more broadly to microfluidics, (3) revised figure 2 to include images from the LCD printer for direct comparison to channels printed from the DLP printer, (4) edited figure 3 to highlight dimensional fidelity comparisons between the two LCD printers, and (5) we have changed the term mSLA printers to LCD printers to use more accessible language that will appeal to a wider audience.

