## Supplementary Information for "Democratizing access to microfluidics: Rapid prototyping of open microchannels with low-cost LCD 3D printers"

Supporting information includes additional experimental details and methods, including a link to a repository of CAD files, raw data, and excel image analysis sheets, LCD printer schematic with pixel alignment depiction of CAD design (PDF), a visualization of how the XY measurements were taken with the dimensional analysis software built-in to our Keyence microscope (PDF), information about optimal printing parameters and settings (PDF), layer thickness calibration chip with SEM photos and measurements (PDF), UV light intensity measurements across the LCD printers’ build plates (PDF), print location across the build plate and measurements of capillaric features (PDF), and a video documenting the performance of capillary domino valves printed on one of our LCD printers (MOV).

### Data Availability

Link to CAD files, raw data (images), and Excel sheets for image analysis: <https://doi.org/10.5281/zenodo.8347134>

### Evaluation of uniform UV LED light intensity methods

Light intensity of the LCD printers was measured with a 405 nm UV light meter (Chitu Systems, Shenzhen, Guangdong, China). The length and width of each LCD screen were divided into four equidistant segments to define 16 equal areas on the LCD screen. Each measured area is 15.2 cm^2^ for the Anycubic Mono X 6K and 7.7 cm^2^ for the Phrozen Sonic Mini 8K. The UV light meter probe was placed on each predefined area and the printer was lit up by pressing Exposure in Tools on the Anycubic Mono X 6K and Vat Cleaning in Tools on the Phrozen Sonic Mini 8K. Each area was lit up for 10 seconds and the highest light intensity within this timeframe was recorded. Measurements were taken for all 16 areas to map the light intensity of the LCD on each 3D printer.

### Designing and evaluating microchips for perpendicularity study methods

Microchannels of various aspect ratios were created to evaluate channel perpendicularity in relation to channel design. The different aspect ratios were 1:9, 1:3, 1:2, and 2:1 and all microchannels were designed for 90° between the side and bottom walls. Each microchip was designed to have N=3 microchannels of different aspect ratios, with a total of 12 microchannels per microchip. The largest height and width values for the microchannels were 160 µm and 344 µm, respectively. The smallest height and width values were 40 µm and 103.2 µm, which were the smallest reproducible values obtained from the XY calibration microchip measurements. All microchannels were measured using the angle measurement tool on the Keyence digital microscope.

### S1: LCD printer overview

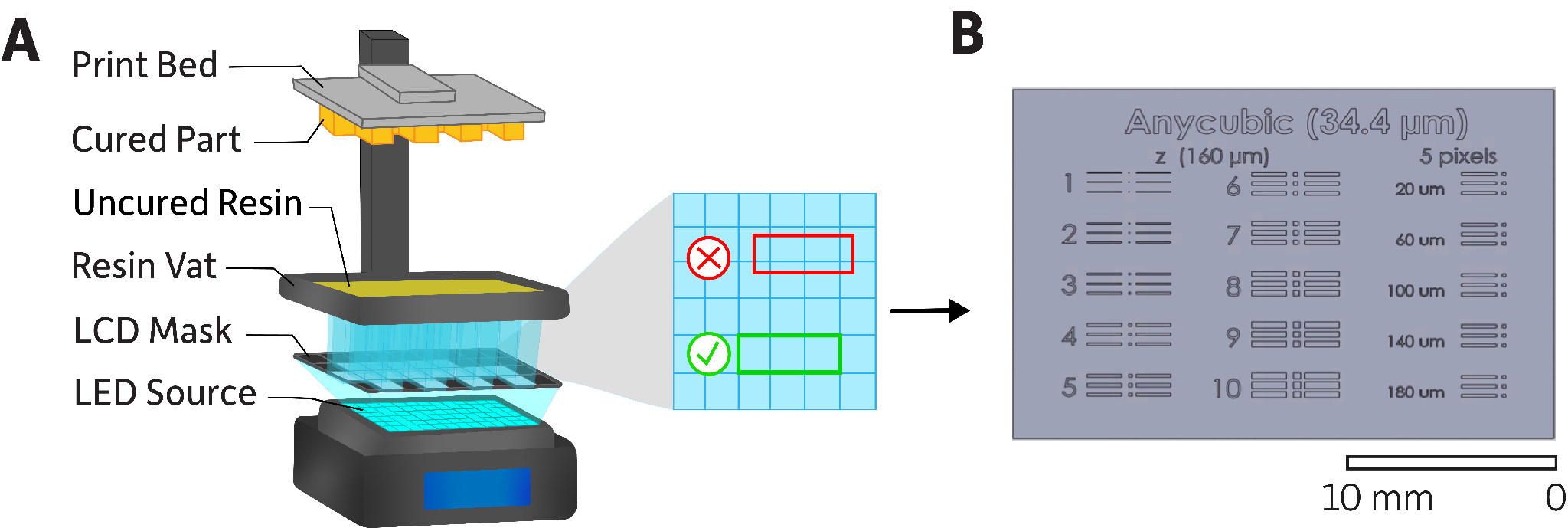

**Supplementary Figure 1: LCD printer schematic with pixel grid alignment and pixel calibration chip**

**(a)** Schematic showing LCD printer components. The LCD mask has a discrete pixel grid. To the right, the pixel grid is depicted with a misaligned feature shown with a red rectangle and a correctly aligned feature shown with a green rectangle. **(b)** CAD design of the XY pixel calibration chip for the LCD printer. The first two columns are channels with fixed height (160 µm) and XY dimensions ranging from 1 to 10 pixels in 1-pixel (34.4 µm) increments. The third column (far right) has features with fixed width (5 pixels) and heights varying from 20 to 180 µm in 40 µm increments.

### S2: Pixel calibration chip XY measurements

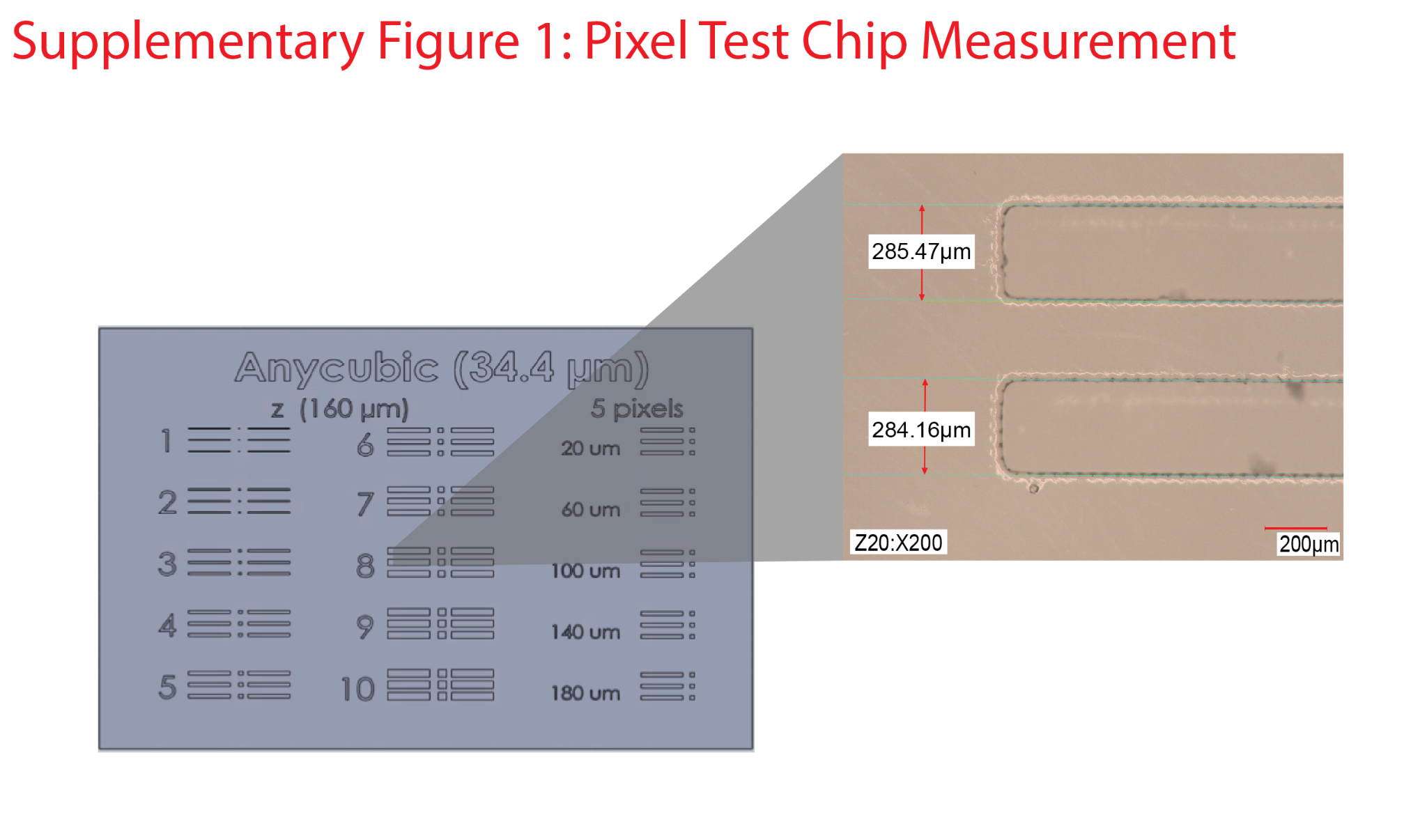

###### **Supplementary Figure 2: Pixel calibration chip prepared for 34.4 µm LCD printer.**

Inset: Photo of 8-pixel channels showing how XY measurements for each channel were taken. It is worth mentioning that the dimensional accuracy measurements can be subjective depending on who measured the variable and proper lighting conditions on the microscopes, as such, there is slight room for variability.

### S3: Initial optimization of print parameters on LCD printers

We systematically varied printing parameters including UV power, exposure time, layer height, retraction speed, off-time, and anti-aliasing to determine the best conditions for printing sub-millimeter, negative features with a minimum overall print time. The manufacturer’s recommended settings for each LCD printer are well suited for printing larger (> 1 mm) features and have not been validated for printing microfluidic features. We found that channel walls were more corrugated as UV power increased on the LCD printer **(Supplementary Figure 3)**. We chose a UV power of 45% that provided straight side walls without excess uncured resin on the microchannel surface.

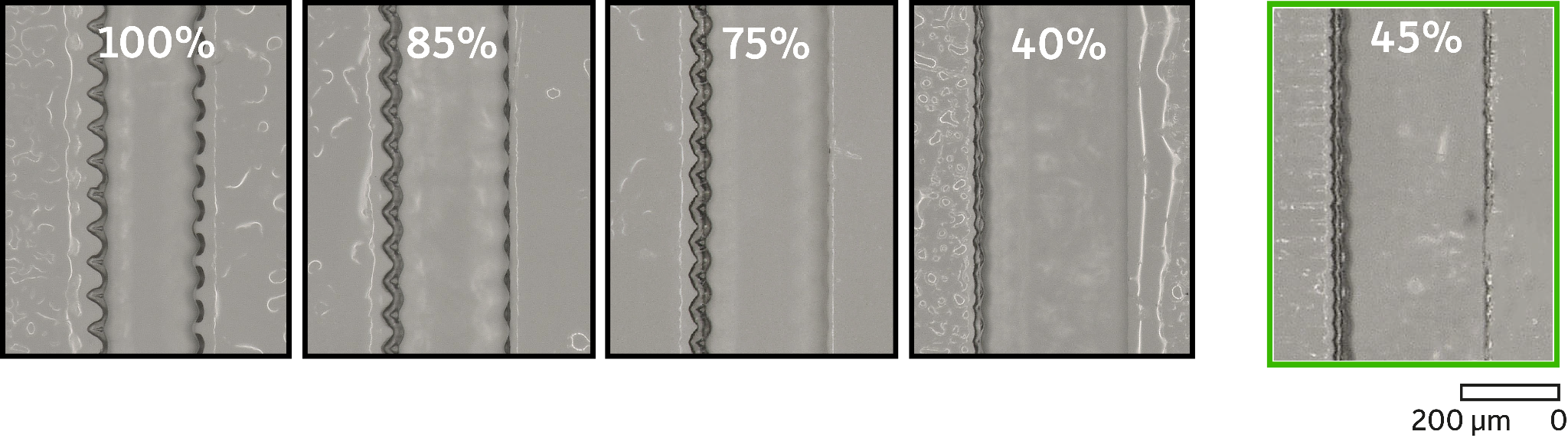

###### **Supplementary Figure 3: Varying UV power on an LCD printer affects the smoothness of microchannel side walls.** We screened UV power ranging from 40% to 100%. We chose a UV power of 45% because the channel sidewalls demonstrated the least amount of corrugation and a minimum amount of uncured resin on the channel surface.

We found that an exposure time of 1.8 seconds produced higher fidelity negative features than the manufacturer’s recommended setting of 2 seconds. Furthermore, increasing the off-time – the time between when the build platform is completely lowered into the resin vat and when the UV light is turned on – from the recommended 0.5 seconds to 1 second improved the quality of our prints. At equivalent layer heights of 20 µm, both the DLP-SLA and 34.4 µm LCD printer produced the same pixel calibration chip in < 34 minutes.

### Table S1: Optimized printing parameters for both LCD printers

###### **Supplementary Table 1: Optimal print parameters for LCD printers.**

|  | **Anycubic Mono X 6K + Water-Wash+ Resin/CADworks resin** | **Phrozen Sonic Mini 8K + CADworks resin** | **Phrozen Sonic Mini 8K + Water-Wash+ Resin** |
| --- | --- | --- | --- |
| **Normal Exposure Time (sec)** | 1.8 | 2.4 | 1.4 |
| **Off time (sec)** | 1 | 1 | 5 |
| **Bottom Exposure Time (sec)** | 23 | 23 | 23 |
| **Bottom Layers** | 6 | 6 | 6 |
| **Layer thickness (mm)** | 0.02 | 0.02 | 0.02 |
| **Normal layer Z lift speed (mm/s)** | 1 | 1 | 1 |
| **Normal layer Z Retract speed (mm/s)** | 1.5 | 1.5 | 1.5 |
| **Anti-alias** | 1 | 1 | 1 |
| **UV Power** | 45% | 100%* | 100%* |

***** UV power was not tunable on the Phrozen Sonic Mini 8K.

### S4: Layer thickness calibration chip Z measurements

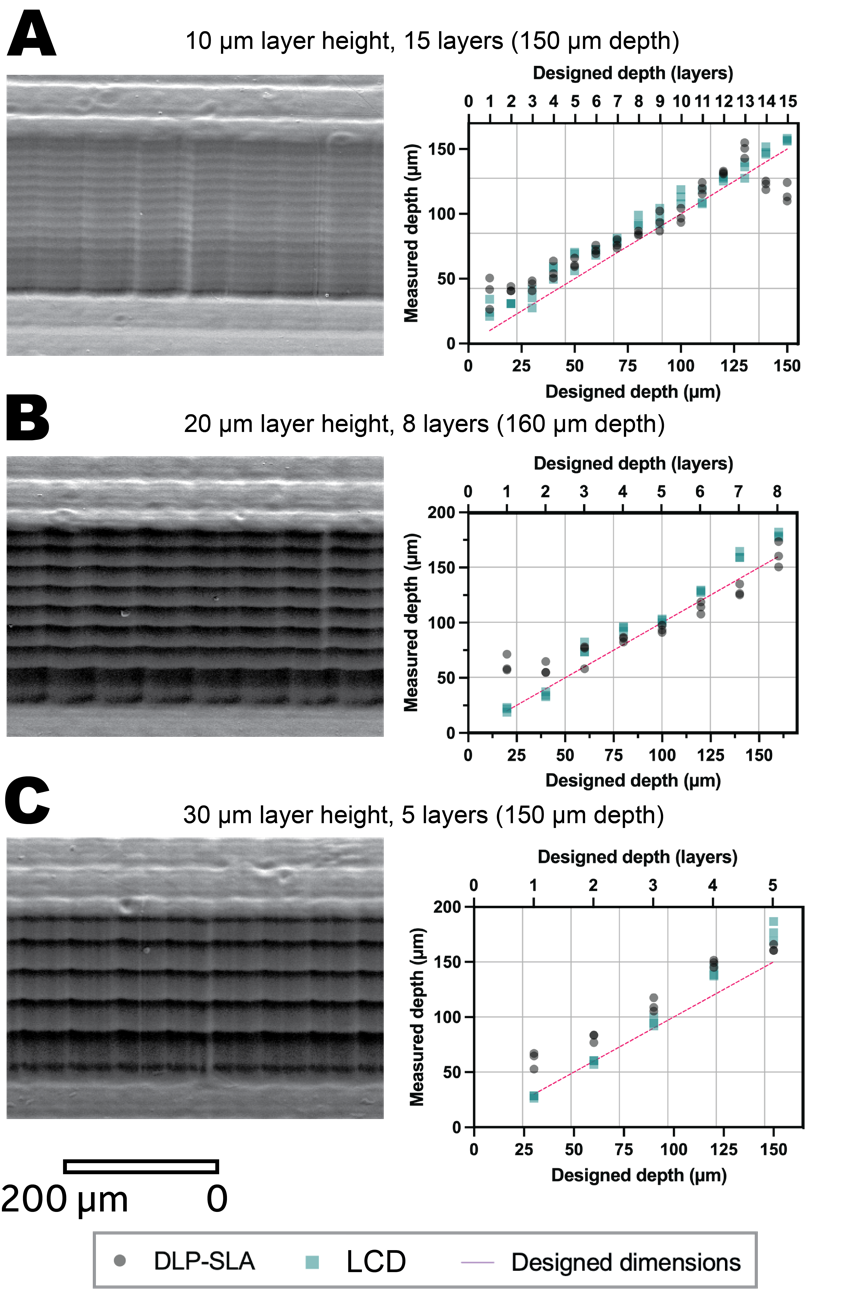

**Supplementary Figure 4: Qualitative and quantitative examination of layer height and feature depth for 3D printed open microchannels.** Accuracy of depth measurements for both the DLP-SLA and LCD printer was comparable across layer heights of 10, 20, and 30 µm. **(a)** Channel printed with a 10 µm layer height and measured depths for different numbers of layers printed, peaking at 15 layers. **(b)** Channel printed with a 20 µm layer height and measured depths for different numbers of layers printed, peaking at 8 layers. **(c)** Channel printed with a 30 µm layer height and measured depths for different numbers of layers printed, peaking at 5 layers.

The lowest documented layer thickness provided by the manufacturers range from 5-10 µm^1,2^, however these values are often incorrect and out of the range of printability. In **Supplementary Figure 4**, as layer thickness is increased from 10 µm to 30 µm the sidewall texture is more pronounced. The 1 layer depths for almost every chip had a substantial CV and deviation from designed values, as such, they were omitted from the following data observations. The highest percent error documented is 23.88% ± 27.99 (**S4A, DLP**) and the smallest is 10.59% ± 8.34 (**S4B, LCD**). While the percent errors remained variable across the different conditions, the CVs remained consistent around ~5% and <= the largest CV of 7.59±4.72 (**S4B, DLP**). Given the variability of the Z-height measurements we recommend printing any critical parameters in the XY dimension given that we can predict the outcome of those measurements with more certainty.

### S5: Printing calibration chips on the 22 µm LCD printer with IPA wash

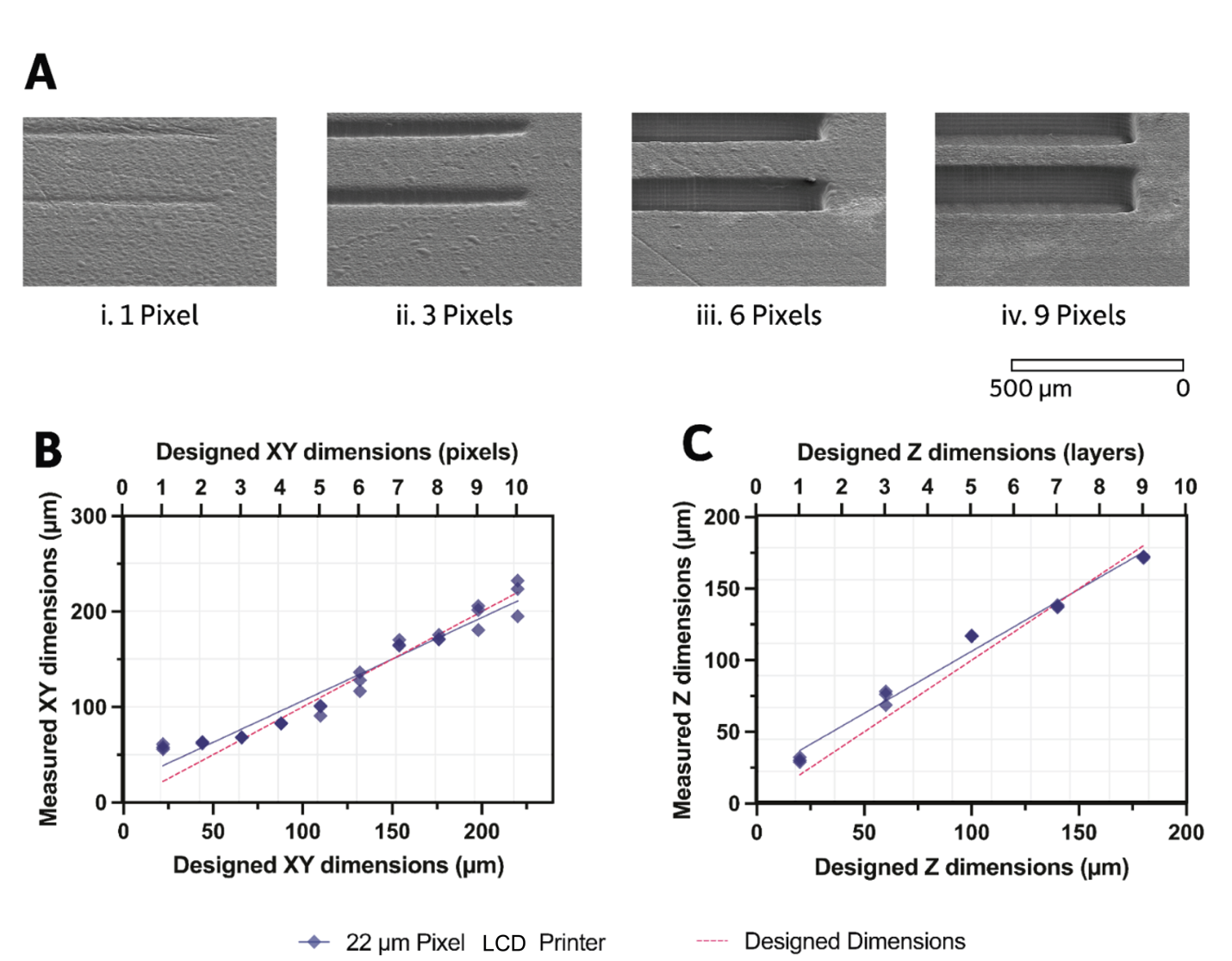

###### **Supplementary Figure 5: 3D printed microchannels with 22 µm pixel LCD printer.**

**(a)** SEM images show that deep microchannels (160 µm height; 13 layers) had high fidelity above 3 pixels in width. **(b)** Microchannels had excellent agreement between designed and measured widths above 3 pixels. **(c)** Measured depths of 3D printed microchannels followed a linear trend with increasing layer height and only slightly overestimated designed dimensions.

### S6: UV light intensity across the build plate of LCD printers

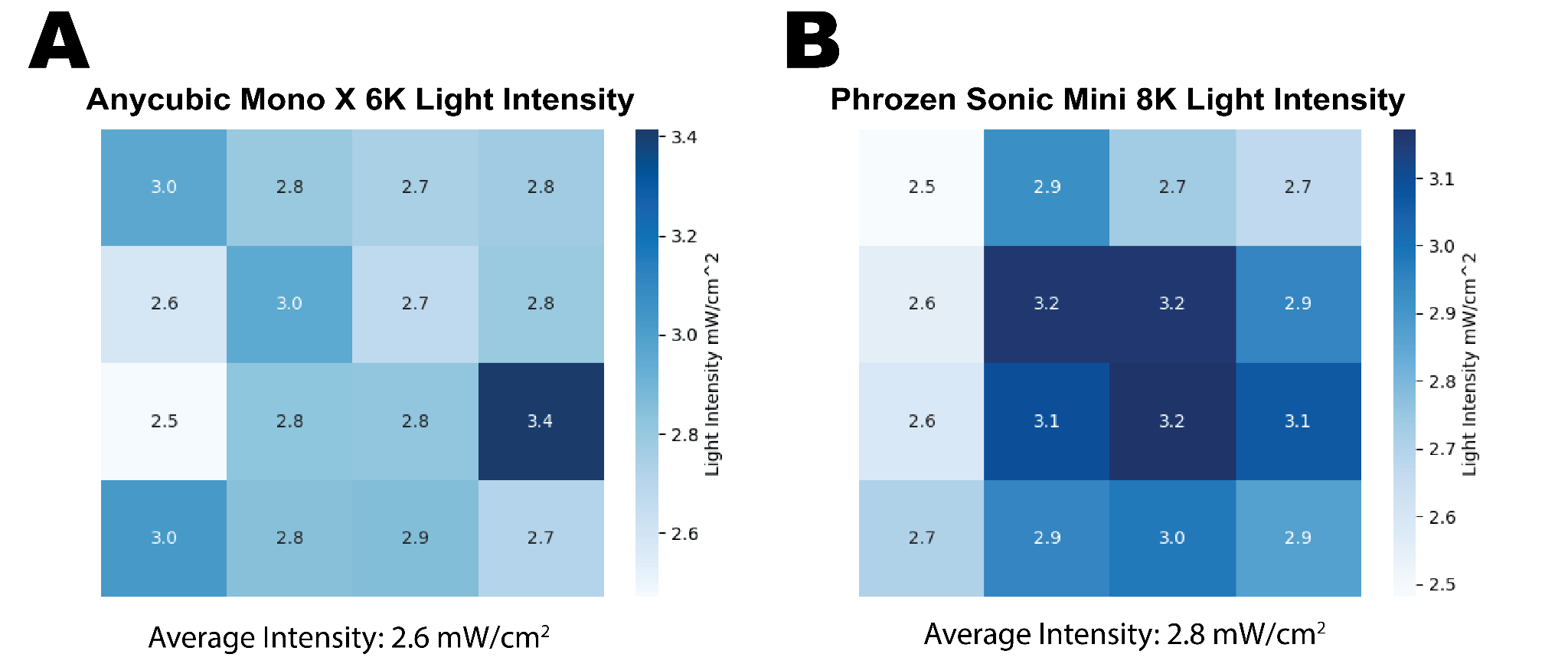

**Supplementary Figure 6: UV light intensity measurements of the Anycubic Mono X 6K and Phrozen Sonic Mini 8K taken across the build area.** **(A)** Light intensity measurements for the Anycubic Mono X 6K were generally uniform, albeit with a few outliers near the perimeter of the build area. **(B)** Light intensity measurements for the Phrozen Sonic Mini 8K are within ±15% of the average intensity.

**Supplementary** **Figure 6A** depicts the light intensity across the Anycubic Mono X 6K build plate with light values representing lower intensities and darker values representing high intensities. The approximate average intensity for the Anycubic at 45% UV power is 2.6 mW/cm^2^. A majority of the values obtained from the light intensity study were higher than 2.6 mW/cm^2^, but within 15% mW/cm^2^ of the average intensity, albeit one lower value situated at 2.5 mW/cm^2^ and one higher outlier situated at 3.4 mW/cm^2^. The values in the center of the build plate were generally uniform with ranges around 2.8 mW/cm^2^, while the areas along the perimeter of the build plate demonstrate more variance.

The average light intensity of the Phrozen Sonic Mini 8K is 2.8 mW/cm^2^. The light intensity values across the build plate **(Supplementary Figure 6B)** are within ± 15% mW/cm^2^ of the average intensity. Similar to the Anycubic, the center of the build plate depicts uniform values, while the perimeter exhibits more variance. The Phrozen has a fixed UV light intensity, while the Anycubic has the ability to modulate UV power; this may affect the accuracy of the light intensity.

### S7: Placement and measurement of capillary domino valve features across the build plate

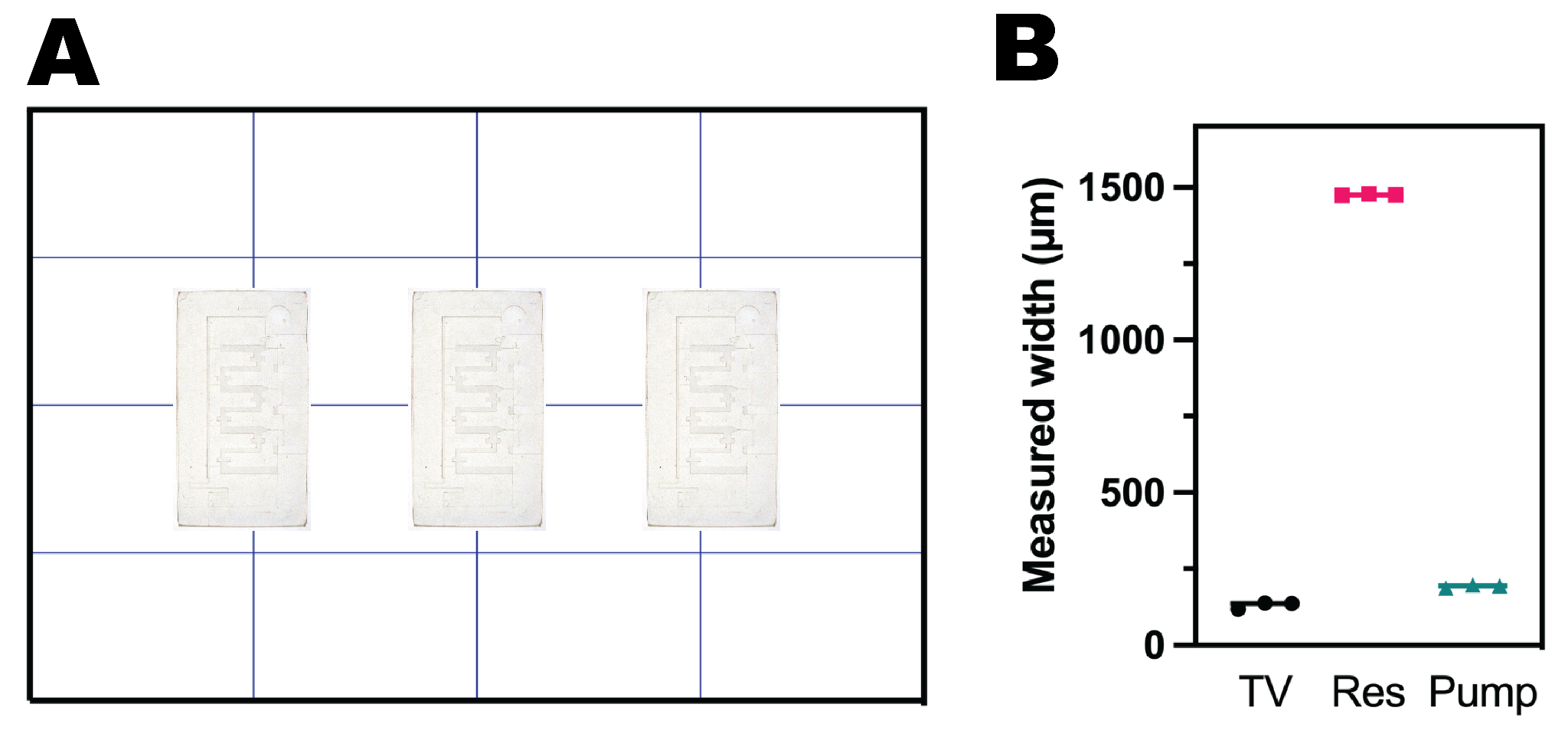
**Supplementary Figure 7: Capillaric circuit locations across the build plate with associated width measurements of small features. (a)** The grid depicts the approximate locations where microchips were printed across the build plate. **(b)** The corresponding graph denotes the measurements of trigger valves, reservoirs, and capillary pumps in the same location across the three different microchips. Regardless of where the microchips were printed on the build plate, they demonstrated minimal variability.
